# A scanning electron microscopy-based screen of leaves of *Solanum pennellii* (ac. LA716) x *Solanum lycopersicum* (cv. M82) introgression lines provides a resource for identification of loci involved in epidermal development in tomato

**DOI:** 10.1101/688077

**Authors:** J. Galdon-Armero, M. L. Arce-Rodriguez, C. Martin

## Abstract

The aerial epidermis of plants plays a major role in their environment interactions, and the development of its cellular components -trichomes, stomata and pavement cells- is still not fully understood. We have performed a detailed screen of the leaf epidermis of two generations of the well-established *Solanum pennellii* ac. LA716 x *Solanum lycopersicum* cv. M82 introgression line (IL) population using a combination of scanning electron microscopy techniques. Quantification of the trichome and stomatal densities in the ILs revealed 18 genomic regions with a low trichome density and 4 ILs with a high stomatal density. We also found ILs with abnormal proportions of different trichome types and aberrant trichome morphologies. This work has led to the identification of new, unexplored genomic regions with roles in trichome and stomatal formation and provides an important dataset for further studies on tomato epidermal development that is publically available to the research community.

## Introduction

The epidermis is the external cell layer of plant organs and, in aerial tissues, consists of three types of specialised cells: trichomes (hairs), stomata and pavement cells. Trichomes are outgrowths (commonly referred to as hairs) which can have different sizes and shapes, and their morphology has been used commonly for taxonomic purposes (Payne, 1978). In *Solanum lycopersicum* (tomato) and related species, trichomes are multicellular and have been classified into seven different types according to size, morphology and metabolic profiles of those bearing glandular heads (Luckwill, 1943, Simmons and Gurr, 2005). Stomata are pores surrounded by two specialised guard cells in which turgor is regulated to control gas exchange between the plant and the atmosphere (Hetherington and Woodward, 2003). Stomata, in contrast to trichomes, have conserved morphology and function in plants (Chater et al., 2017). Pavement cells are relatively unspecialised epidermal cells which ensure an adequate patterning of trichomes and stomata on the leaf surface. In tomato, pavement cells have a characteristic undulated shape, like pieces of a jig-saw puzzle, a shape which is not uncommon in other dicotyledonous plants (Vőfély et al., 2019). These different cell types emerge from a single cell layer, the protodermis, and therefore are developmentally linked, as suggested by studies in tobacco and tomato (Glover et al., 1998, Glover, 2000). The importance of correct spacing of epidermal cell types and the potentially limited size of the pool of protodermal cells implies cross-talk in the regulation of determination of different cell fates in the epidermis.

From a developmental point of view, most studies of trichome formation have focused on the model plant *Arabidopsis thaliana*, which produces only one type of unicellular, non-glandular trichome, contributing to the establishment of a detailed model for their initiation and development at the cellular and molecular levels (Pattanaik et al., 2014, Szymanski et al., 1998). The transcriptional regulation of trichome initiation in *A. thaliana* involves the formation of a MYB-BHLH-WD40 (MBW) complex which induces trichome formation. The main MYB transcription factor forming part of this complex is GLABROUS1 (GL1) (Larkin et al., 1994), although MYB23 and MYB82 can perform the same function redundantly (Kirik et al., 2005, Liang et al., 2014). Three BHLH factors can form part of this MBW complex, GLABRA3 (GL3) (Payne et al., 2000) and ENHANCER OF GLABRA3 (EGL3) (Zhang et al., 2003), and TRANSPARENT-TESTA8 (TT8) plays the same role in leaf margins (Maes et al., 2008). Additionally, a WD40 factor has been identified as part of the complex, TRANSPARENT TESTA GLABRA1 (TTG1) (Walker et al., 1999). This complex activates the expression of downstream genes necessary for trichome development. One such target is the WRKY transcription factor encoded by *TRANSPARENT TESTA GLABRA 2 (TTG2)*. TTG2 has been suggested to be recruited to the MBW complex itself through interaction with the WD40 protein TTG1 (Pesch et al., 2014; Lloyd et al., 2017). Alternatively, TTG1 and TTG2 may interact downstream of the MBW complex to narrow the target genes responding to its transcriptional regulation (Lloyd et al., 2017) Among other targets, *GLABRA2* (*GL2*) encodes an HD-Zip transcription factor essential for the correct morphogenesis of the mature trichomes (Szymanski et al., 1998). Furthermore, the MBW complex can be activated by a hierarchical cascade of C2H2 zinc finger transcription factors, including GLABROUS INFLORESCENCE STEM proteins (GIS, GIS2 and GIS3) and ZINC FINGER PROTEIN 5, 6 and 8 (Gan et al., 2007, Sun et al., 2015, Zhou et al., 2011, Zhou et al., 2013, Gan et al., 2006). In contrast, negative regulation of trichome initiation involves the expression of small R3 MYB transcription factors in the trichome initial including TRYPTICON (TRY), CRAPICE (CPC), ENHANCER OF TRYPTICON AND CAPRICE1/2/3 (ETC1/2/3) and TRICHOMLESS1/2 (TCL1/2). These R3 MYB proteins can move to neighbouring cells and compete with GL1 in the formation of the MBW complex (Wester et al., 2009, Wada et al., 1997, Schnittger et al., 1999, Kirik et al., 2004b, Kirik et al., 2004a). However, this model does not apply to tobacco, tomato or related species (Serna and Martin, 2006), where trichome formation is controlled by different regulatory proteins (Lloyd et al., 2017).

In tomato, the focus of research has been on glandular trichomes and the metabolites they secrete (Schilmiller et al., 2008, McDowell et al., 2011). A number of different transcription factors involved in trichome formation have been identified. SlMX1 (a MIXTA transcription factor) was shown to control trichome initiation in tomato, while also regulating cuticle deposition and carotenoid content in fruit (Ewas et al., 2016). MIXTA transcription factors are known regulators of trichome initiation in several species such as *Artemisia annua* (Yan et al., 2018), cotton (Wu et al., 2018) or *Populus* (Plett et al., 2010). Moreover, two HD-ZIP transcription factors have been identified as regulators of trichome development in tomato, Woolly (Yang et al., 2011) and CUTIN DEFICIENT2 (CD2) (Nadakuduti et al., 2012). Woolly controls trichome initiation and specifically the morphogenesis of long, glandular type I trichomes (Yang et al., 2011) and does so by forming a complex with a small cyclin, SlCycB2, which regulates the mitotic divisions in multicellular trichomes (Gao et al., 2017). CD2 regulates cuticle deposition and the formation of glandular type VI trichomes (Nadakuduti et al., 2012), and the development of this trichome type is also regulated by the basic helix-loop-helix transcription factor SlMYC1 (Xu et al., 2018). A C2H2 zinc finger transcription factor, HAIR, has been identified as a regulator of the formation of both type I and type VI trichomes and may control trichome initiation (Chang et al., 2018). Finally, the tomato homolog of TRYPTICHON (SlTRY) is functionally equivalent to TRY when ectopically expressed in *A. thaliana*, but its native function in tomato remains uncharacterised (Tominaga-Wada et al., 2013). However, current understanding of the regulation of trichome initiation and development in tomato is much more limited than for Arabidopsis.

The use of wild tomato species as a source of genetic variation has resulted in the identification of important quantitative trait loci (QTLs) for many traits (Rick and Chetelat, 1995), and has been undertaken traditionally by screening of near-isogenic introgression lines (ILs), generated by successive backcrossing of the offspring of a cross between a wild relative species and a cultivated crop to the cultivated parent (Eshed and Zamir, 1995). The most widely used IL population in tomato is the *S. pennellii* ac. LA716 x *S. lycopersicum* cv. M82 IL population, which has been extensively curated and genotyped precisely (Chitwood et al., 2013). Comprehensive analyses of this IL population has led to the identification of loci of interest including tolerance to pathogens (Smart et al., 2007, Sharlach et al., 2013), abiotic stress (Frary et al., 2011, Rigano et al., 2014), primary metabolism (Magnum et al., 2018) and morphogenesis (Chitwood et al., 2013, Ron et al., 2013). With respect to trichomes, extensive work on characterising trichome secretion in the ILs has been undertaken, revealing QTLs involved in the synthesis of acyl sugars and terpenoids (Schilmiller et al., 2010), and a visual assessment of trichome phenotypes in the IL population aided in the identification of HAIR as a regulator of trichome formation (Chang et al., 2018). However, the aerial epidermis of the ILs has not been characterised fully and we still lack detailed understanding of the degree of variability present in the population. Differences in trichome and stomatal densities and trichome types reported for *S. lycopersicum* and *S. pennellii* (Simmons and Gurr, 2005, McDowell et al., 2011, Heichel and Anagnostakis, 1978a) support the use of the *S. pennellii* ac. LA716 x *S. lycopersicum* cv. M82 ILs as a platform for investigating trichome development in tomato.

We have performed a comprehensive analysis of the leaf epidermis of two generations of the *S. pennellii* ac. LA716 x *S. lycopersicum* cv. M82 introgression lines (ILs) by a combination of Scanning Electron Microscopy (SEM) techniques. The outputs of this study constitute an important resource for further research into cellular development and led to the identification of unexplored genomic regions associated to the determination of stomatal and trichome density in leaves, as well as trichome morphogenesis.

## Results

### -Identification of genomic regions involved in the determination of leaf trichome density

We evaluated the adaxial leaf epidermis of fully expanded first true leaves of the IL population over two generations, and we identified specific ILs consistently showing differences for specific parameters over the two generations. We measured trichome density in the parental lines *S. lycopersicum* cv. M82 and *S. pennellii* ac. LA716 (Fig 1A) as well as the ILs over two generations (Fig. 1B and S1). Seedlings were grown for 3-4 weeks until the first true leaves were fully expanded. For each leaf sample 8-15 micrographs of the adaxial epidermis of the same area of a leaflet of the one of the first true leaves were prepared. Three to four seedlings were scanned for each IL in each generation.

**Figure 1.**
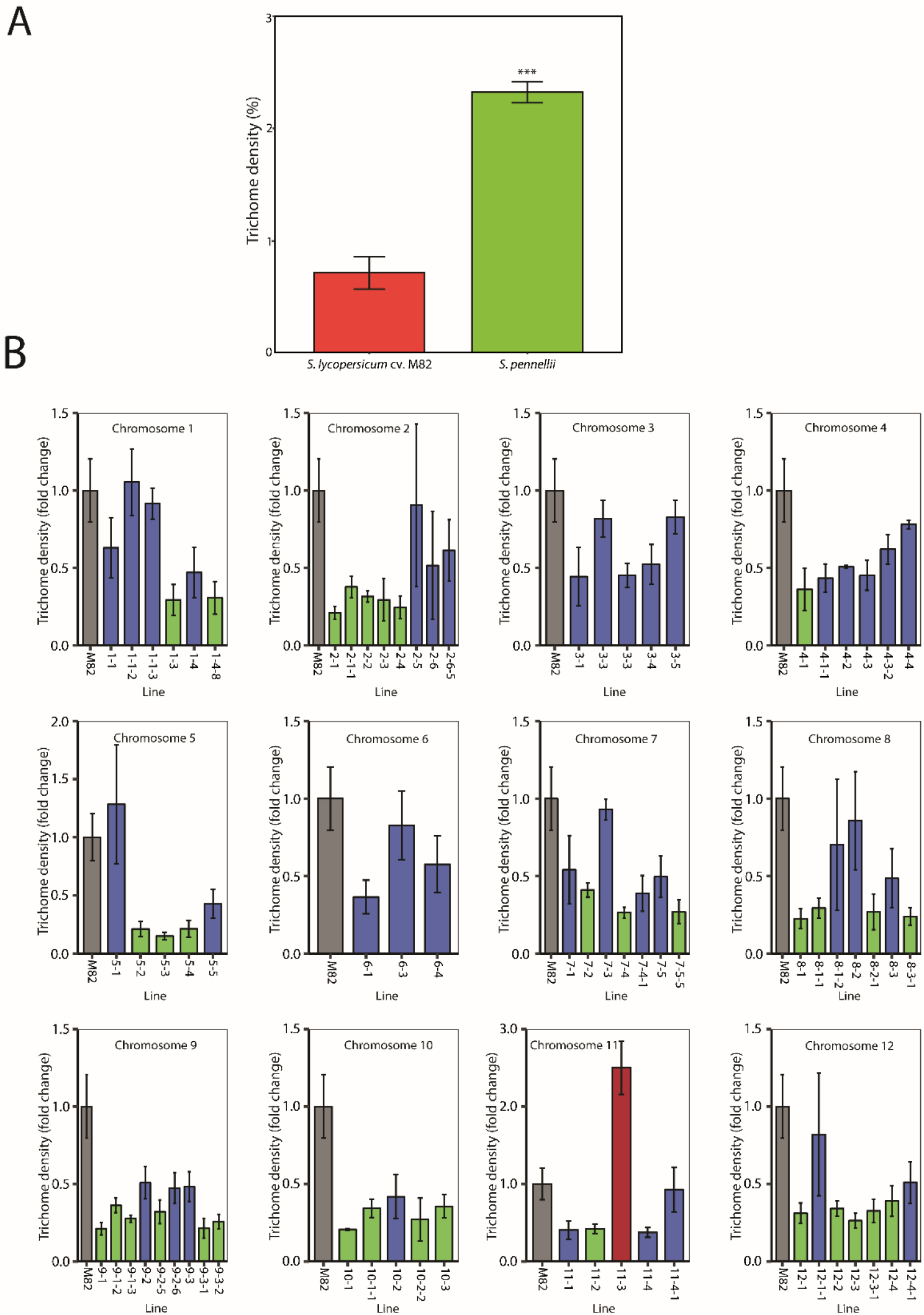
Trichome densities of the *S. pennellii* (ac. LA716) x *S. lycopersicum* (cv. M82) ILs. **A)** Trichome density of the first fully expanded true leaf of *S. lycopersicum* cv. M82 (red bar) and *S. pennellii* ac. LA716 (green bar). Values are expressed as mean ± SEM (n=3). Stars indicate significant differences (p-value<0.01) according to t-tests. **B**) Trichome density of the first generation of ILs, grouped according to the chromosomal location of the introgressed *S. pennellii* genomic region. Values are mean ± SEM (n=3-4) of the relative value of trichome density compared to M82 values (grey bar). Significant differences were determined using t-tests between the value for each IL and the value for M82. Green bars indicate ILs with significantly lower trichome densities than M82 (p-value<0.05) and red bars indicate ILs with significantly higher trichome densities than M82 (p-value<0.05). Blue bars indicate other ILs.

We measured the trichome and stomatal density of *S. pennellii* ac. LA716 and *S. lycopersicum* cv. M82 parents on the adaxial surface of leaves and found it was three-fold higher in *S. pennellii* ac. LA716 compared to *S. lycopersicum* cv. M82 (Fig. 1A and 2A). This is in agreement with previous reports on trichomes in *Solanum* species (Simmons and Gurr, 2005) but contrasts with previous reports of stomatal density in *S. pennellii*, where stomatal density in *S. pennellii* was reported to be lower than in *S. lycopersicum* (Chitwood et al., 2013, Heichel and Anagnostakis, 1978b). These differences supported the use of *S. pennellii* as a donor species to study epidermal development in *Solanum*.

**Figure 2.**
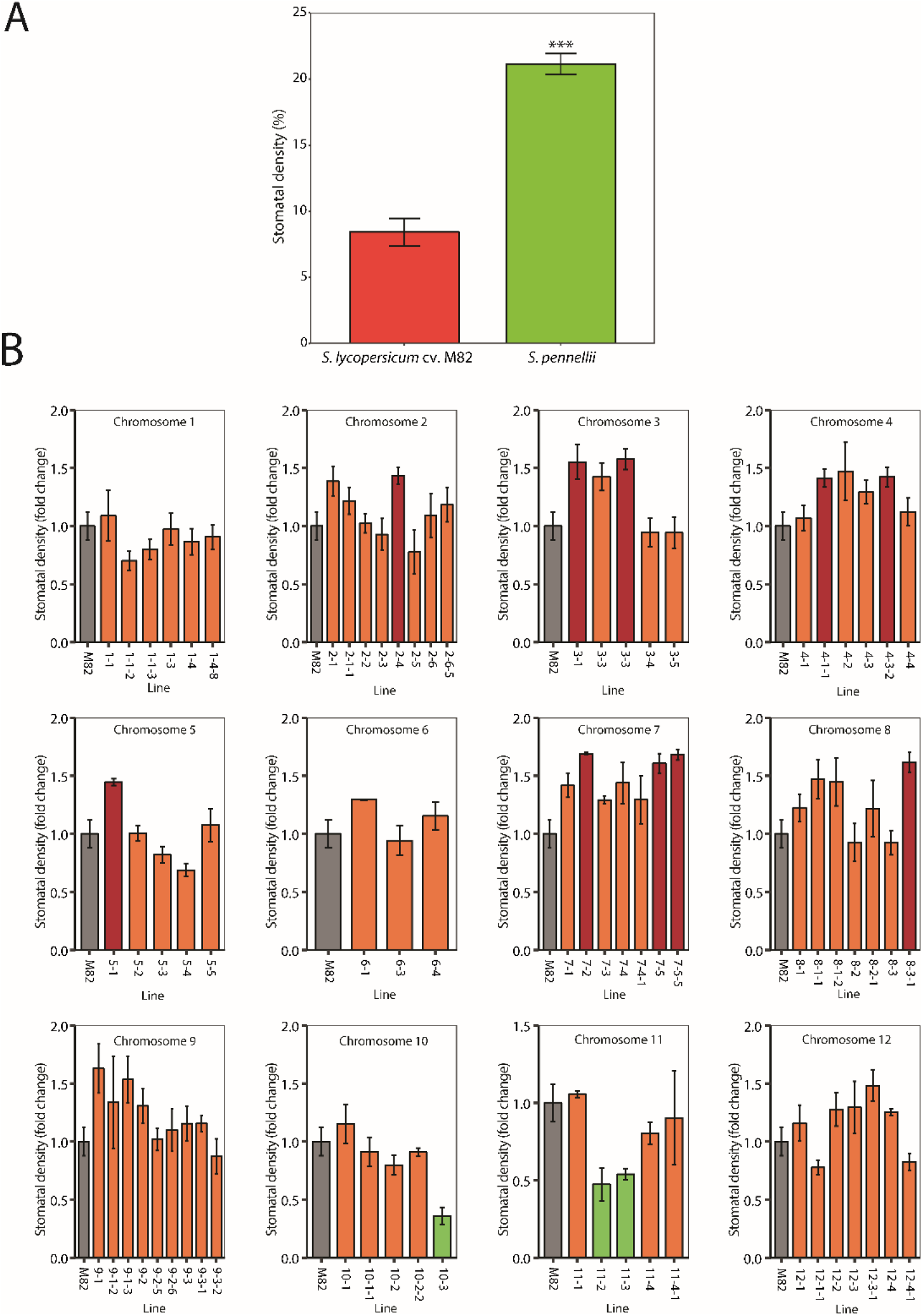
Stomatal density of the *S. pennellii* (ac. LA716) x *S. lycopersicum* (cv. M82) ILs. **A)** Stomatal density of the first fully expanded true leaf of *S. lycopersicum* cv. M82 (red bar) and *S. pennellii* ac. LA716 (green bar). Values are expressed as mean ± SEM (n=3). Stars indicate significant differences (p-value<0.01) according to a t-test. **B)** Stomatal density of the first generation of ILs, grouped according to the chromosomal location of the introgressed *S. pennellii* genomic region. Values show the mean ± SEM (n=3-4) of the relative value of stomatal density compared to M82 values (grey bar). Significant differences were determined using t-tests between the value for each IL and the value for M82. Green bars indicate ILs with a significantly lower stomatal density than M82 (p-value<0.05) and red bars indicate ILs with a significantly higher stomatal density than M82 (p-value<0.05). Orange bars indicate any other ILs.

We observed 19 ILs with low trichome density over the two generations: specifically ILs 2-1-1, 2-2 and 2-3 on chromosome 2; IL 4-1 in chromosome 4; ILs 5-3 and 5-4 on chromosome 5, IL 7-2 on chromosome 7; ILs 8-1, 8-1-1 and 8-2-1 on chromosome 8; ILs 9-1-3, 9-3-1 and 9-3-2 on chromosome 9; ILs 10-1 and 10-3 on chromosome 10; ILs 11-2 on chromosome 11; and ILs 12-1, 12-3 and 12-3-1 on chromosome 12. IL 11-3 had a significantly higher trichome density only in the first generation, but in the second generation, although not significantly different from M82, it averaged the highest value for trichome density.

### -Identification of genomic regions involved in the determination of leaf stomatal density

We measured the stomatal density of the parental lines *S. lycopersicum* cv. M82 and *S. pennellii* ac. LA716 (Fig 2A) and over two generations of the IL population (Fig. 2B and S2). We identified four ILs with high stomatal density. These were: IL 5-1 on chromosome 5; ILs 7-2 and 7-5 on chromosome 7 and IL 8-3-1 on chromosome 8. No IL showed consistently lower stomatal density than M82.

### -Identification of genomic regions involved in the determination of trichome types

We classified trichomes according to the categories established by Luckwill et al., (1943) over two generations. Trichomes were extensively damaged by the use of chemical fixation and critical point drying of the samples from the first generation (Fig. S3), and therefore we focused our analysis on the data from the second generation (Fig. 3). In M82, the main trichome type is type I/IV, accounting for 63% of the total. The other types of trichome found in *S. lycopersicum* were observed in smaller percentages (5-10%). We used the distribution in M82 as a standard for comparison with the rest of the ILs. For most lines, type I/IV trichomes were the most abundant, although ILs 2-1, 3-3, 4-3-2,8-1- 1 and 8-2-1 showed substantial reductions in this type of trichome. This reduction in type I/IV trichomes was compensated by an increase in non-glandular type V trichomes in ILs 2-1 and 3-3. For 21 of the ILs, we did not observe any type V trichomes on the adaxial surface of the first true leaf. Type VI trichomes were generally observed at low frequency, but type VI trichomes were the most common type in IL 8-1-1, 8-2-1 and 8-3-1. A total of 6 ILs, ILs 2-4, 2-6-5, 4-1, 4-1-1, 5-5 and 9-3 showed no type VI trichomes in the assessed tissue. Type VII trichomes were the rarest and absent on the first true leaves of 31 of the ILs. Importantly however, the absence of any one type of trichome from the adaxial epidermis of the first true leaf does not imply a complete absence of this trichome type in other tissues.

**Figure 3.**
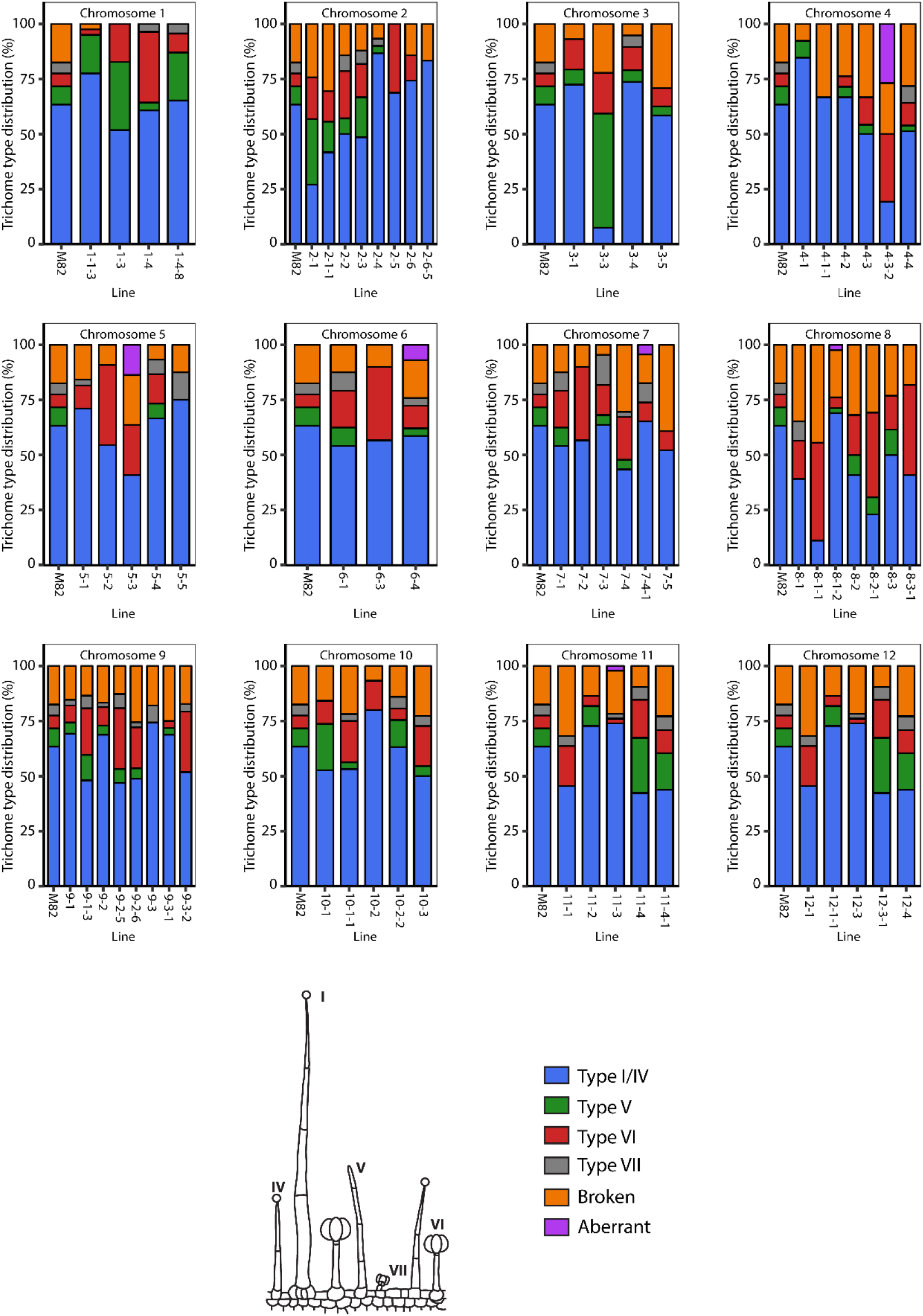
Trichome type distribution of the *S. pennellii* (ac. LA716) x *S. lycopersicum* (cv. M82) ILs. Lines are classified according to the chromosomal region introgressed from *S. pennellii* and each bar represents an IL, and the height of each colour section represents the proportion in percentage of each type of trichome. Blue represents type I/IV trichomes; green represents type V trichomes; red represents type VI trichomes; grey represents type VII trichomes, orange represents damaged trichomes and purple represents aberrant trichomes. A schematic representation of each type of trichome is displayed in the legend. Trichomes are classified in different types according to (Luckwill, 1943).

### -Identification of genomic regions involved in trichome morphogenesis and spatial patterning

For each line under study, trichomes showing aberrant morphology were recorded (Fig. 4 and S4). The most common type of aberrant trichome found in the population consisted of two swollen cells emerging from a type I-like multicellular base. This type of trichome was observed in ILs 4-3-2, 5-3, 6-4, 7-5, 8-1-2 and 11-3 (Fig. 4B-G). Trichomes with aberrant division patterns were also found in ILs 1-1-2, 5-1 and 7-4-1 (Fig. S4A-C). In IL 10-2, we observed another type of aberrant trichome, consisting of branched, multicellular, non-glandular trichomes (Fig. S4D). In every IL with aberrant trichomes, the aberrant forms always appeared alongside wild type trichomes of the same type in both generations (Fig. 3 and S3).

**Figure 4.**
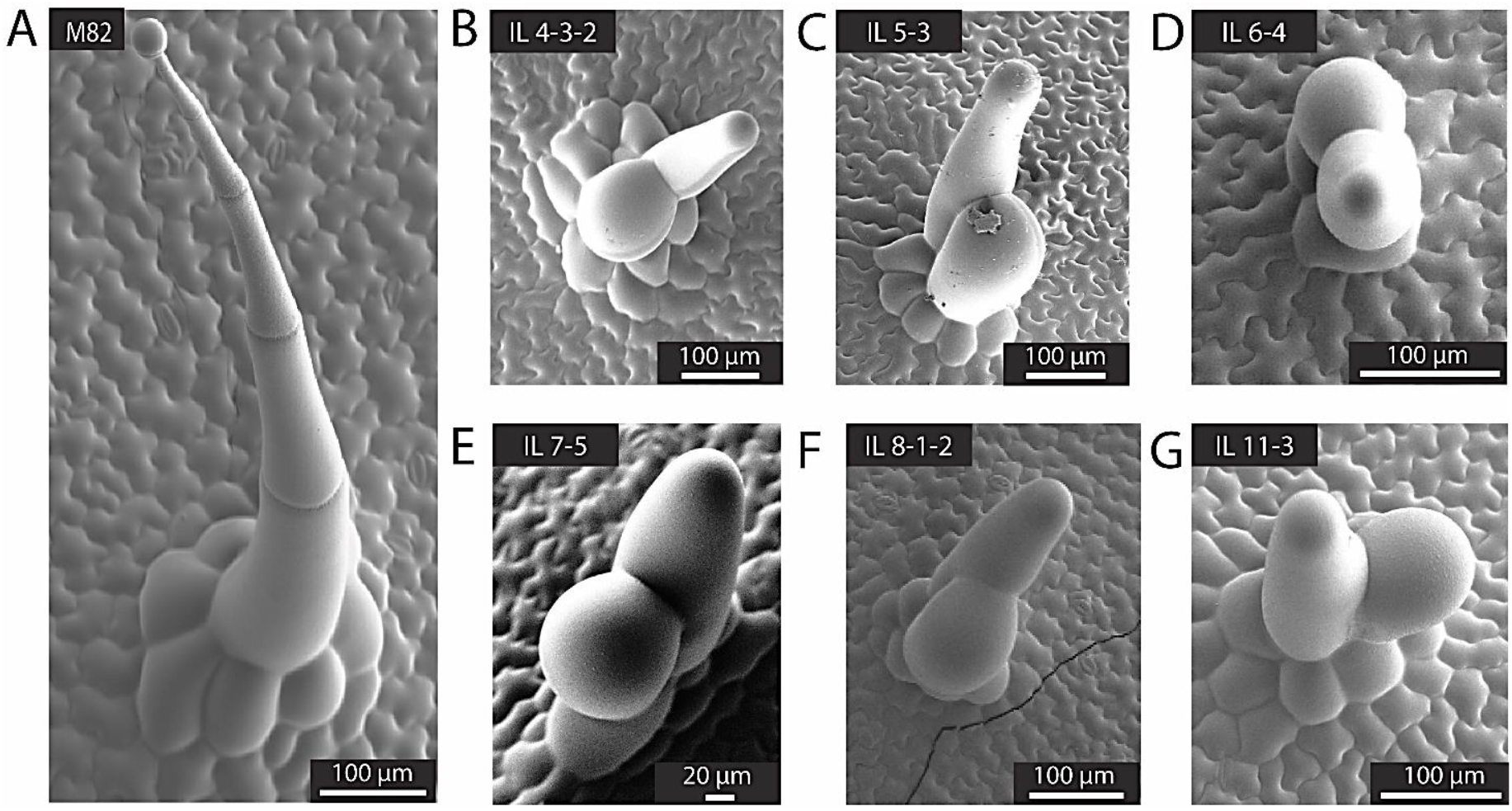
Aberrant trichome morphologies in the IL population. **A)** SEM micrograph of a representative type I trichome on the leaf surface of M82. Micrographs of representative aberrant trichomes are shown for ILs **B**) 4-3-2, **C**) 5-3, **D**) 6-4, **E**) 7-5, **F**) 8-1-2, **G**) 11-3, all sharing a similar swollen and forked appearance. Scale bars are shown in each micrograph.

Finally, we observed unusual clusters of trichomes on the adaxial surface of leaves in ILs 2-5 and 2-6 (Fig. 5). In young leaves, trichomes in M82 are equally distributed on the leaf surface and they are oriented in the same direction (Fig. 5A). However, in IL 2-5 trichomes were clustered in groups of up to 4 trichomes and they were randomly oriented (Fig. 5B). In mature, fully expanded leaves, we could still observe clusters of two trichomes in ILs 2-5 and 2-6, and these observations were made for glandular and non-glandular trichomes (Fig. 5). Trichomes in these clusters were found oriented in all possible directions respectively to each other.

**Figure 5.**
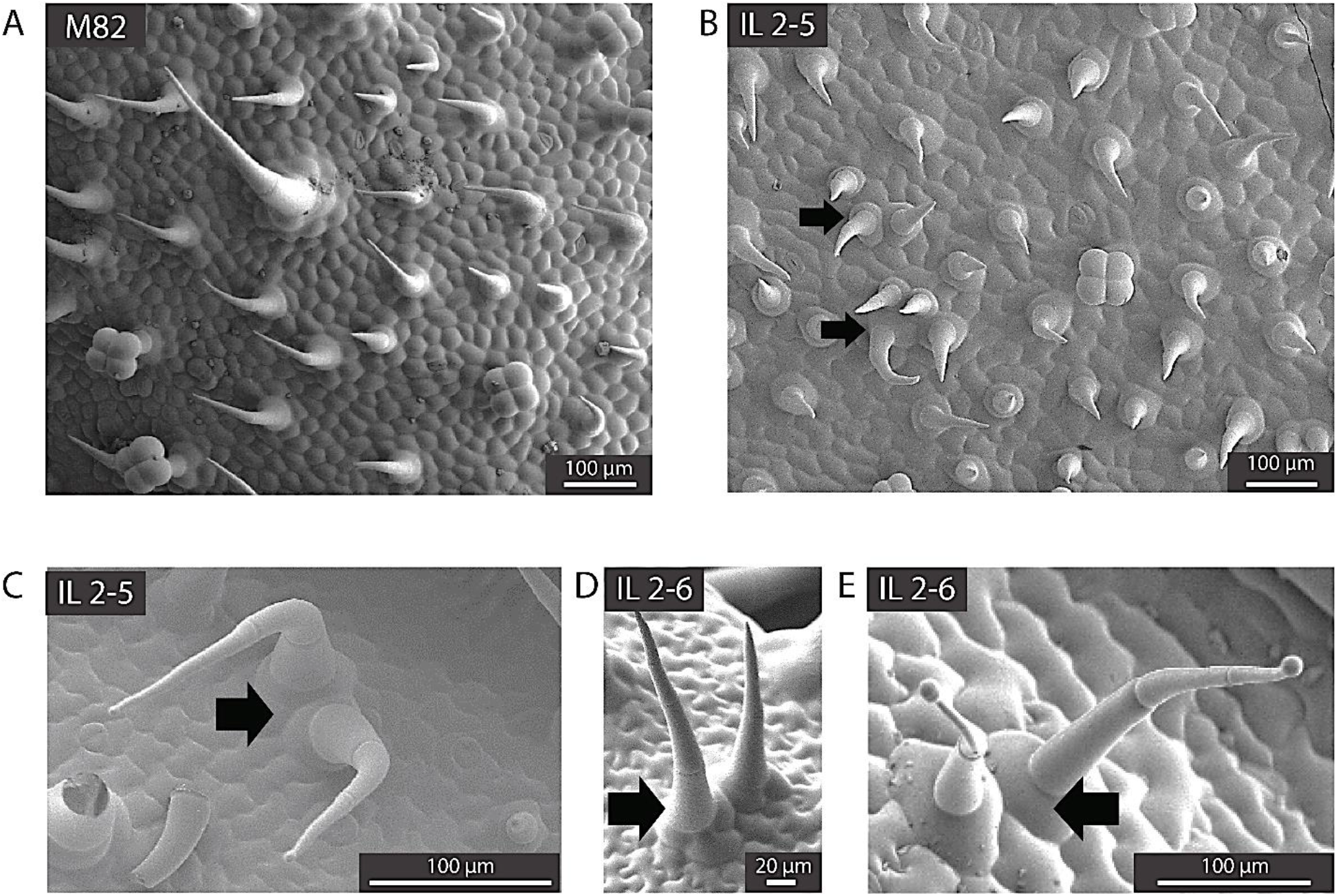
Abnormal trichome clusters in the IL population. **A**) SEM micrograph of the adaxial surface of a young leaf of M82. **B**) SEM micrograph of the adaxial surface of a young leaf of IL 2-5. **C**) SEM micrograph of a cluster of two type IV trichomes in IL 2-5. **D**) SEM micrograph of a cluster of two type V trichomes in IL 2-6. **E**) SEM micrograph of a cluster of two type IV trichomes in IL 2-6. Scale bars are shown for each micrograph. Black arrows indicate clusters of trichomes.

## Discussion

### -Natural variation in epidermal development in the *S. lycopersicum* cv. M82 x *S. pennellii* ac. LA716 ILs

Through comparison to *S. pennellii* ac. LA716 and to *S. lycopersicum* cv. M82 (Fig. 1A and 2A) we identified 18 ILs that showed consistent phenotypes over two generations, suggesting stable genetic components regulating trichome density in these ILs. Interestingly, most of the ILs analysed showed lower trichome density than M82, and these differences were consistent in both generations of the IL population (Fig. 1B and S1B). This was unexpected given the values observed for the parents (Fig. 1A). However, the complexity of trichome development, which involves cell wall expansion, cell division and differentiation (Glover, 2000) and the negative and positive control exerted by different regulatory factors (Tian et al., 2017) could well be impaired by small changes in the activity or expression of genes introgressed in the ILs. Some known genes involved in trichome development map to the genomic regions corresponding to the 18 ILs of interest. An aquaporin-like gene (Solyc08g066840) has been proposed as a candidate gene for the *dialytic* (*dl*) mutant phenotype (Chang et al., 2016), which maps to the IL 8-2-1 region. However, this mutant was not compromised in trichome initiation but rather in trichome development, displaying aberrant, forked trichomes on its leaf surface. The auxin-responsive factor 3 (SlARF3) plays an important role in the development of epidermis in tomato, and when silenced, trichome density was reduced (Zhang et al., 2015). SlARF3 maps to the region covered by IL 2-3. The HD-Zip transcription factor responsible for the *Woolly* (Solyc02g080260) mutant phenotype, with increased trichome density (Yang et al., 2011), also maps to the IL 2-3 region of chromosome 2. These genes are potential candidates for determining the phenotype observed in this line (Fig. 1 and S1). The Woolly transcription factor interacts with a type-B cyclin (SlCycB2, Solyc10g083140), essential for division of multicellular trichomes, which causes a dramatic reduction in trichome density when silenced in RNAi lines (Gao et al., 2017). This gene maps to the region covered by IL 10-3, and might also be responsible for the low trichome density in this line. Further analysis of these ILs will be necessary to determine whether other factors might be involved in the determination of trichome density.

We assessed trichomes on the adaxial surface of the first true leaf of two generations of the ILs, using chemical fixation coupled with room temperature SEM for the first generation (Fig. S3) and cryo-scanning electron microscopy (SEM) for the second generation (Fig. 3). Trichomes were classified according to (Luckwill, 1943), although we grouped type I and type IV in the same group based on their similar morphology and metabolic profiles as proposed by (McDowell et al., 2011). In a similar fashion, non-glandular trichomes are traditionally classified as type II, III or V depending on their length, but for comparative purposes we classified them all as type V (Fig. 3 and S3). The most abundant trichome type in M82 and most ILs was type I/IV (Fig. 3, in contrast to previous reports, that ranked type VI trichomes as the most common glandular trichomes (Bergau et al., 2015). However, the density of type I trichomes has been reported to be a juvenility trait, being very high in the first leaf - the leaf we assessed (Fig. 3 and S3) (Vendemiatti et al., 2017). We observed five lines with a low proportion of type I trichomes (Fig. 3). In the case of ILs 2-1 and 3-3, this was compensated by an increase in the percentage of type V trichomes, in contrast to the observations in Vendemiatti et al. (2017), making these good candidate lines for further research into the control of the juvenile phase in vegetative development of tomato. In the case of ILs 8-1-1 and 8-2-1, the observed reduction in type I trichomes was accompanied by a reduction in trichome density (Fig. 1 and 3). This suggests that the pathway controlling the formation of type I trichomes may be compromised in these lines, while other types of trichome might not be affected. Both lines also showed an increased proportion of type VI trichomes. This could indicate that different types of trichome are controlled by different regulatory mechanisms and that these mechanisms might be interlinked to compensate for alterations in the distribution between trichome types, in agreement with previous reports in tomato (Li et al., 2004, Yang et al., 2011) and tobacco (Payne et al., 1999).

Type VI trichomes were relatively unabundant in the sampled leaves, but absent only in 6 ILs of the population. IL 4-1 had no type VI trichomes, and this line also had low overall trichome density (Fig. 1 and 3). Similar to the ILs on chromosome 8, this low trichome density could be due to a specific reduction in type VI trichomes. However it is important to note that, although we could not find type VI trichomes on the adaxial surface of the first true leaves, they were found on stems and major veins, suggesting that IL 4-1 influences tissue-specific regulation of epidermal development, which has been reported in other species such as *A. thaliana* (Schnittger et al., 1998) or *Antirrhinum majus* (Glover et al., 1998). Type VII trichomes were rare in the interveinal leaf tissue of all lines, resulting in our finding none in almost half of the lines (Fig. 3). This type of trichome was, however, commonly found on major and minor veins, indicating that their number is higher in epidermis overlying vascular tissues.

We found several ILs with aberrant trichome morphologies (Fig. 4 and S4) and unusual epidermal patterning (Fig. 5). The most common aberrant morphology consisted of a multicellular base similar to that found in type I trichomes, with two swollen cells emerging from this base (Fig. 4). These structures are reminiscent of the aberrant type I trichomes observed in the *odourless-2* mutant (Kang et al., 2010) and the *dialytic* mutant (Chang et al., 2016). The swollen appearance of these trichomes is similar to that observed in the *hairless* mutant (Kang et al., 2016), which presents a truncated version of the *SPECIALLY-RAC1 ASSOCIATED* (*SRA1*) gene and is consequently compromised in the organisation of the actin cytoskeleton in trichomes, or the *inquieta* mutant, which harbours an altered copy of the *ACTIN-RELATED PROTEIN 2/3 COMPLEX SUBUNIT 2A*(*ARPC2A*) (Jeong et al., 2017). It is possible that the observed phenotype might be due to alterations in the process of cell elongation or cytoskeleton development in trichomes. In fact, the *ARPC2A* gene maps to the IL 11-3 region of tomato (Fig. 4G). The *ACTIN-RELATED PROTEIN 3* (*ARP3*) and *ACTIN-RELATED PROTEIN 2/3 COMPLEX SUBUNIT 3* (*ARPC3*) genes encode proteins that are part of the complex required for actin filament formation (Goley and Welch, 2006), and map to the IL 4-3-2 and IL 7-5 regions respectively (Fig. 4B and E). All of these aberrant trichomes appear amid wild-type looking trichomes (Fig. 3 and S3) pointing towards phenotypes caused by small changes in expression or functionality of *S. pennellii* alleles of the genes rather than gain- or loss-of-function differences between the two species. We also observed branched, multicellular, non-glandular trichomes in IL 10-2 which lacked a multicellular base (Fig. S4D). This structure resembles the type V-like trichomes observed in SlCycB2 or Woolly RNAi lines (Yang et al., 2011). In fact, SlCycB2 maps to the region delimited by this IL, indicating that the *S. pennellii* allele of the gene might have reduced activity compared to the alleles in M82. Finally, we observed an unusual clustering of trichomes in ILs 2-5 and 2-6 (Fig. 5). Trichomes are evenly distributed over the leaf surface in tomato (Fig. 5A), although the mechanisms by which this patterning is determined are unclear. In *A. thaliana*, trichome patterning is mediated by small MYB transcription factors that inhibit trichome initiation in cells adjacent to newly formed trichomes (Hauser, 2014), but the mechanism in tomato and related species is not yet understood (Tominaga-Wada et al., 2013). These ILs provide useful tools to gain new insights into cell fate determination in the epidermis of tomato leaves.

We also measured the stomatal density for each line in both generations of the ILs (Fig. 2 and S2). We identified 4 ILs with a consistently high stomatal density in both generations. Interestingly, most of the ILs showed a higher stomatal density than the parental line M82 (Fig. 2 and S2). In previous studies a lower stomatal density in *S. pennellii* compared to *S. lycopersicum* has been reported (Heichel and Anagnostakis, 1978, Chitwood et al., 2013), although our study showed a stomatal density significantly higher in *S. pennellii* than in the cultivated parent (Fig. 2A). These differences were the opposite to those observed for trichomes, where most ILs had a lower trichome density than M82 (Fig. 1 and S1). These observations suggested a possible developmental link between the two epidermal structures, perhaps involving an early commitment of cell fate to trichome formation that prevents stomatal formation thereafter as described for tobacco (Glover et al., 1998) and tomato under drought conditions (Galdon-Armero et al., 2018). We analysed the genomic regions covered by the 4 ILs with consistent high stomatal density to determine if any known regulators of stomatal development (Pillitteri and Dong, 2013) mapped to them. We could not identify any known regulatory genes in these introgressed regions, suggesting that novel genes involved in the determination of stomatal density might be responsible for the observed phenotypes.

## Methods

### Plant material

We grew two generations of the *S. pennellii* ac. LA716 x *S. lycopersicum* cv. M82 introgression lines (ILs) (Eshed and Zamir, 1995) to assess epidermal cells on the adaxial leaf epidermis. The first generation consisted of 74 ILs grown under the same conditions, but all seeds were sown simultaneously in October 2016. The second generation consisted of 67 ILs, which were grown under greenhouse condition at the John Innes Centre, with an average temperature of 20-22 °C. These lines were grown successively from October 2015 to February 2016, with approximately 6 ILs phenotyped per week. For both generations, plants were grown for 4-weeks, until the first true leaf was fully expanded. For each line, 3-4 plants in each generation were phenotyped.

### Scanning Electron Microscopy

Trichome phenotypes were assessed using a Zeiss Supra 55 VP SEM (Zeiss, Germany). Two different ways of sample fixation were used to preserve the structure of trichomes and other epidermal structures. For cryo-fixation, plant samples were glued to a cryo-stage and then submerged in liquid nitrogen in a vacuum-generating environment to reduce frosting of water vapour. The samples were then introduced in the microscope preparation chamber, where any frost was removed by sublimation at −95 °C for 3.5 min. The samples were then sputter-coated with platinum for 150 s and transferred to the main microscope chamber, at −125 °C. Imaging took place in the main chamber, where the electron beam was active, and the secondary and back-scattered electrons could be perceived by the detectors. This fixation protocol is destructive, and samples cannot be stored for further imaging, but ensures integrity of most trichomes. Cryo-fixation was used to screen the second generation of ILs.

Chemical fixation was used to screen the first generation of ILs. Chemical fixation of the samples was achieved by vacuum infiltrating them with a glutaraldehyde 2.5% cacodylate solution and dehydrated through an ethanol series. Samples were dried in a Leica CPD300 critical-point dryer (Leica Microsystems, Germany), where water was replaced by liquid CO2 and then evaporated at the critical point for CO2, removing all the liquid without damaging the structures of the sample. Dried samples were glued to a SEM stub and gold-coated before imaging in the main chamber of the microscope, at high vacuum and room temperature conditions.

### Quantification of epidermal structures

For each leaf sample, 8-15 micrographs of 0.3 mm^2^ were generated. In every case, the same leaf (first fully expanded leaf) and the same part of each leaflet (intervein space close to the central vein) was used for assessment of stomatal, trichome and pavement cell numbers. These three cell types were manually quantified in micrographs at a relatively low magnification (x600) using ImageJ v. 1.49 (National Institutes of Health, USA). Trichome and stomatal densities were calculated as percentage of total epidermal cells and expressed as fold change of each line with respect to the M82 values obtained in the corresponding generation. Trichomes were classified in different groups according to Luckwill (1943), except that type I and type IV trichomes were grouped together under type I (McDowell et al., 2011) and all non-glandular trichomes were classified as type V. For trichome density values, all trichomes were considered together. Trichome-to-stomata ratios were calculated by dividing trichome density by stomatal density. Aberrant trichome morphologies were recorded and SEM pictures were taken at the most appropriate magnification. All micrographs used in this study are available in the BioStudies database (http://www.ebi.ac.uk/biostudies) (McEntyre et al., 2015) under accession number S-BSST262.

### Statistical analysis

To assess differences in trichome and stomatal density in the IL population, we performed t-tests to compare the value of each individual line to the value of M82 for the corresponding generation. For the first generation, where values were more consistent, a p-value cut-off of 0.05 was used. For the second generation, where cell density values were more extreme and variable, a p-value cut-off of 0.005 was used. For the trichome type distributions, we made qualitative observations without determining significance. The analyses were performed using R software (ver. 3.2.2; R Core Team, Vienna, Austria).

## Acknowledgements

We acknowledge the financial support of a Rotation Studentship from the John Innes Foundation for JGA, the Institute Strategic Programs: Understanding and Exploiting Plant and Microbial Secondary Metabolism (BB/J004596/1) and Molecules from Nature (BB/P012523/1) from the UK Biotechnology and Biological Sciences Research Council (BBSRC) to JIC.

