## Supplementary Figures 1, 2 3 and 4 for "A scanning electron microscopy-based screen of leaves of *Solanum pennellii* (ac. LA716) x *Solanum lycopersicum* (cv. M82) introgression lines provides a resource for identification of loci involved in epidermal development in tomato"

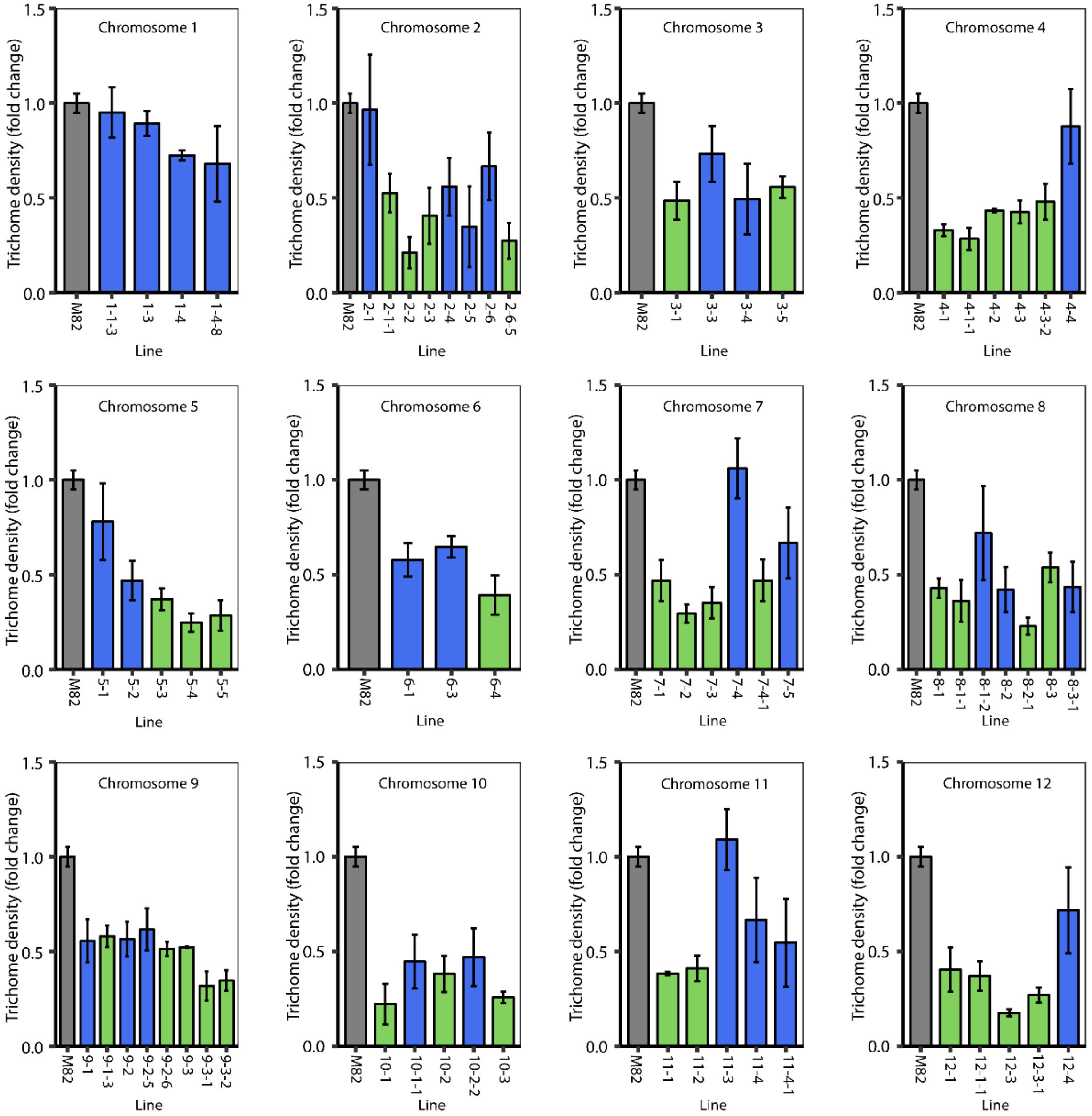


**Supplementary figure 1. Trichome density of the second generation of the *S. pennellii* (ac. LA716) x *S. lycopersicum* cv. M82 ILs.** Trichome density of the second generation of ILs, grouped according to the chromosomal location of the introgressed *S. pennellii* genomic region. Values are mean ± SEM (n=3-4) of the relative value of trichome density compared to M82 values (grey bar). Significant differences were determined using t-tests between the value for each IL and the value for M82. Green bars indicate ILs with a significantly lower trichome density than M82 (p-value<0.005). Blue bars show other ILs.

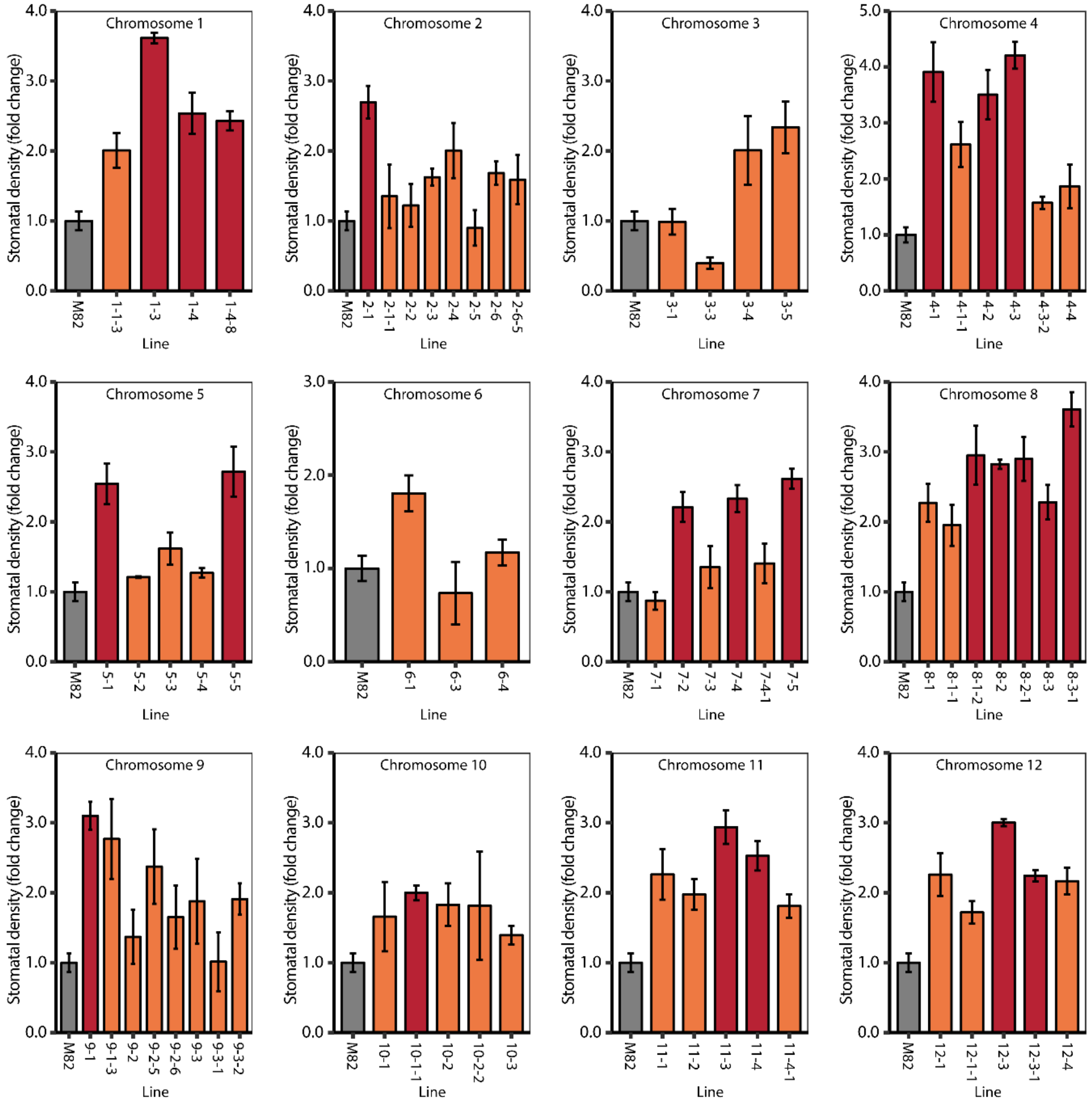


**Supplementary figure 2. Stomatal density of the second generation of the *S. pennellii* (ac. LA716) x *S. lycopersicum* cv. M82 ILs.** Stomatal density of the second generation of ILs, grouped according to the chromosomal location of the introgressed *S. pennellii* genomic region. Values are mean ± SEM (n=3 or 4) of the relative value of trichome density compared to M82 values (grey bars). Significant differences were determined using t-tests between the value for each IL and the value for M82. Red bars indicate ILs with significantly higher stomatal density than M82 (p-value<0.005). Orange bars indicate other ILs.

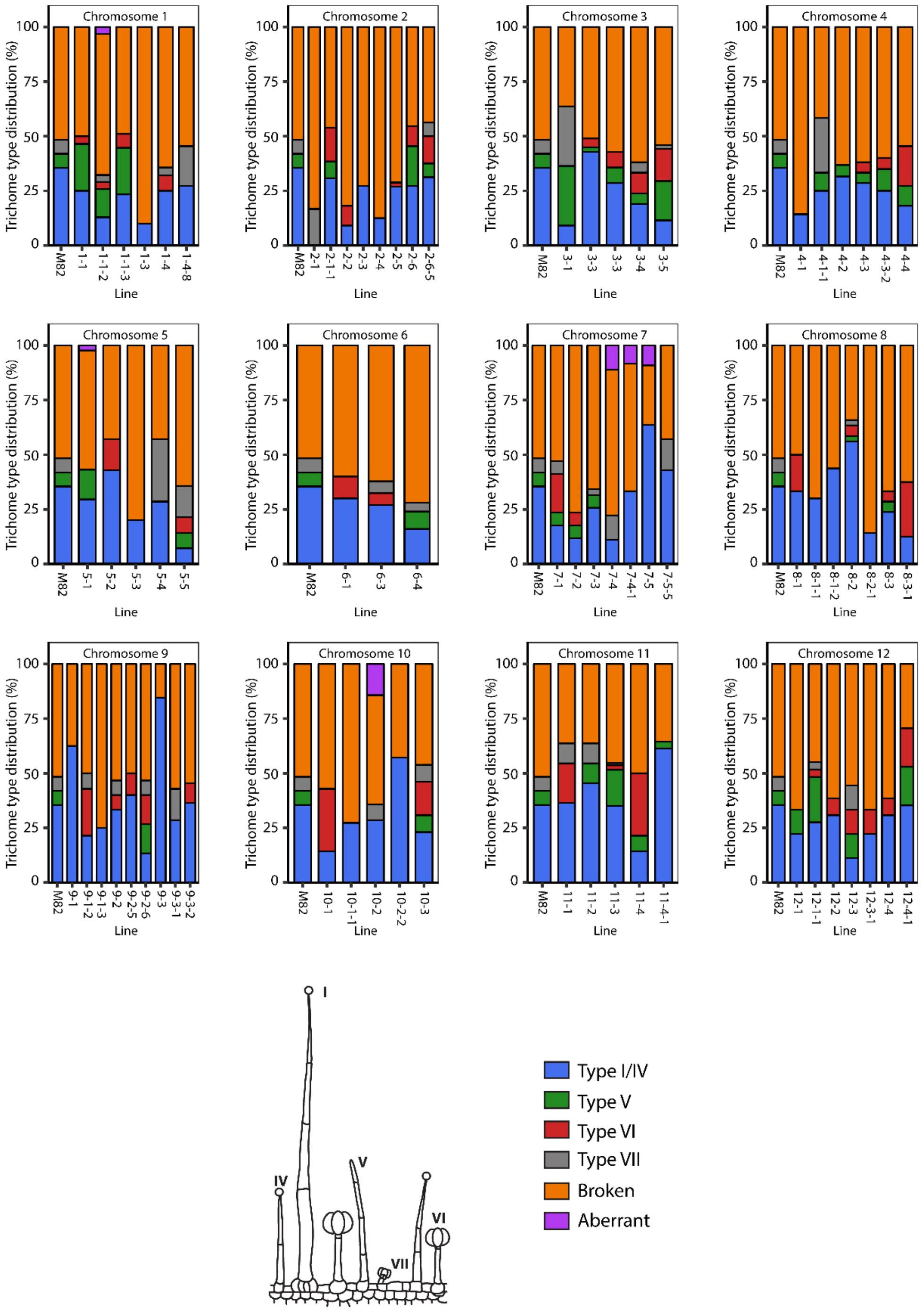


**Supplementary figure 3.-Trichome type distribution of the first generation of *S. pennellii* (ac. LA716) x *S. lycopersicum* cv. M82 ILs.** Lines are classified according to the chromosomal region introgressed from *S. pennellii* and each bar represents an IL, and the height of each colour section represents the proportion in percentage of each type of trichome. Blue represents type I/IV trichomes; green represents type V trichomes; red represents type VI trichomes; grey represents type VII trichomes, orange represents damaged trichomes and purple represents aberrant trichomes. A schematic representation of each type of trichome is displayed in the legend. Trichomes are classified in different types according to Luckwill (1943).

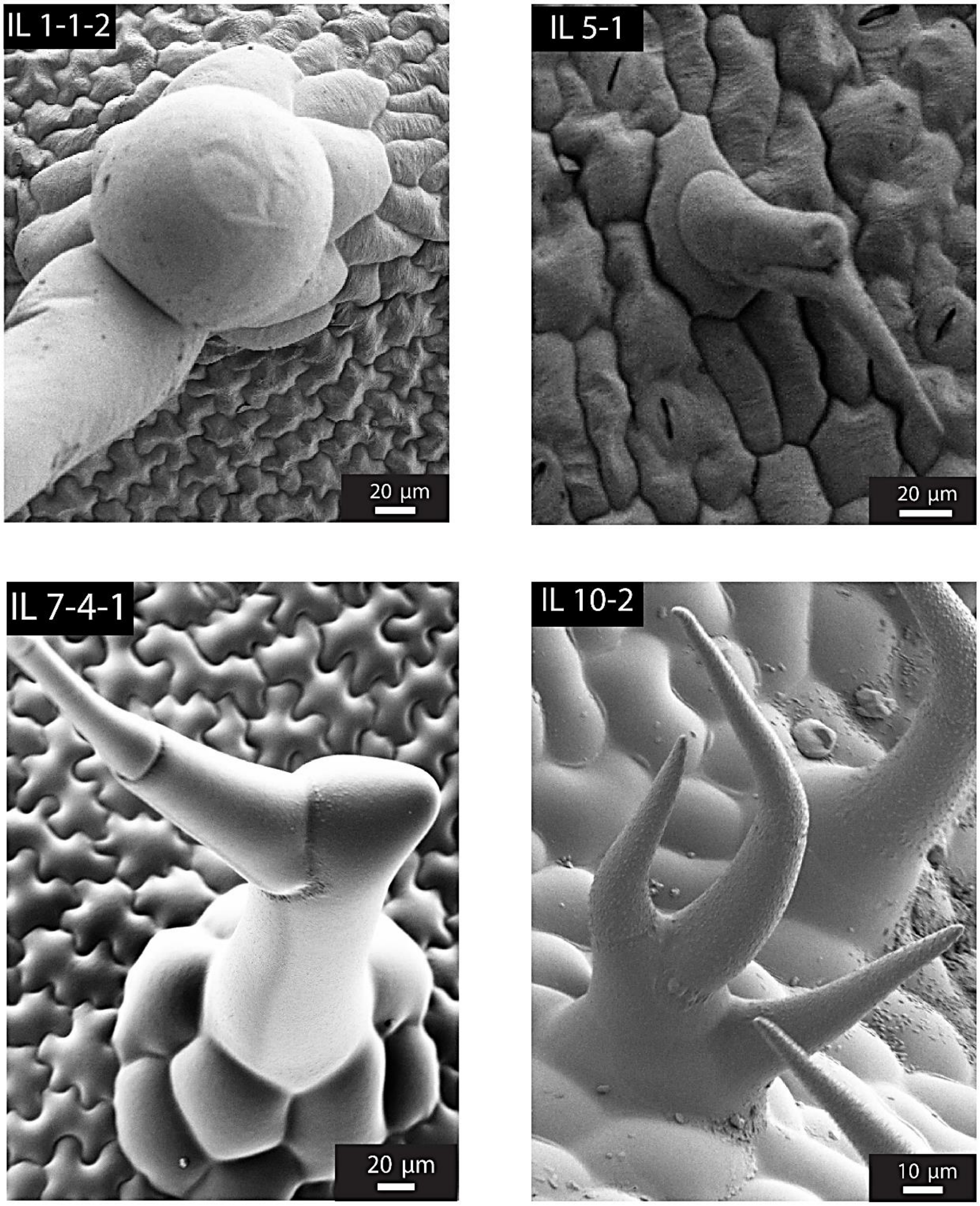


**Supplementary figure 4. Aberrant trichomes found in the IL population. A**) SEM micrograph of an aberrant trichome in IL 1-1-2. **B**) SEM micrograph of an aberrant trichome in IL 5-1. **C**) SEM micrograph of an aberrant trichome found in IL 7-4-1. **D**) SEM micrograph of an aberrant trichome found in IL 10-2. Scale bars are indicated in each micrograph.
